# Ecological Network Metrics: Opportunities for Synthesis

**DOI:** 10.1101/125781

**Authors:** Matthew K. Lau, Stuart R. Borrett, Benjamin Baiser, Nicholas J. Gotelli, Aaron M. Ellison

## Abstract

Network ecology provides a systems basis for approaching ecological questions, such as factors that influence biological diversity, the role of particular species or particular traits in structuring ecosystems, and long-term ecological dynamics (e.g., stability). Whereas the introduction of network theory has enabled ecologists to quantify not only the degree, but also the architecture of ecological complexity, these advances have come at the cost of introducing new challenges, including new theoretical concepts and metrics, and increased data complexity and computational intensity. Synthesizing recent developments in the network ecology literature, we point to several potential solutions to these issues: integrating network metrics and their terminology across sub-disciplines; benchmarking new network algorithms and models to increase mechanistic understanding; and improving tools for sharing ecological network research, in particular “model” data provenance, to increase the reproducibility of network models and analyses. We propose that applying these solutions will aid in synthesizing ecological subdisciplines and allied fields by improving the accessibility of network methods and models.

## Introduction

Interactions are at the heart of ecology and drive many of its key questions. What are the roles of species interactions in ecological systems? When and why is biological diversity important? What factors influence the long-term dynamics of ecosystems? These are all questions with a long history in ecology (Cherrett 1989; Council 2003; Lubchenco et al. 1991; Sutherland et al. 2013) that are not addressed in isolation. Points of intersection include the relationship between diversity and stability (May 2001, 2006); the identity and role of species that are the main drivers of community structure (e.g., keystone species, Paine 1966), ecosystem engineers (Jones et al. 1994), or foundation species (Dayton 1972; Ellison et al. 2005); and the causes and consequences of introducing new species into existing assemblages (Baiser et al. 2008; Simberloff and Holle 1999). Furthermore, “systems thinking” has been a persistent thread throughout the history of ecology (Margalef 1963; Odum and Pinkerton 1955; Patten 1978; Patten and Auble 1981; Ulanowicz 1986), dating back at least to Darwin’s *Origin of Species* in his famous pondering of an entangled bank (Bascompte and Jordano 2014; Golley 1993). The application of network theory has provided a formal, mathematical framework to approach systems (Bascompte and Jordano 2014; Proulx et al. 2005) and led to the development of network ecology (Borrett et al. 2014; Patten and Witkamp 1967; Poisot et al. 2016*b*).

Network ecology can be defined as the use of network models and analyses to investigate the structure, function, and evolution of ecological systems at many scales and levels of organization (Borrett et al. 2012; Eklöf et al. 2012). The influx of network thinking throughout ecology, and ecology’s contribution to the development of network science highlights the assertion that “networks are everywhere” (Lima 2011). And, as one would expect, the field has grown rapidly, from 1% of the primary ecological literature in 1991 to over 6% in 2017 (Fig. 1A). Some examples include: applying network theory to population dynamics and spread of infectious diseases (May 2006); description and analysis of networks of proteins in adult organisms (Stumpf et al. 2007) or during development (Hollenberg 2007); expanding classical food webs to include parasites and non-trophic interactions (Ings et al. 2009; Kéfi et al. 2012); investigating animal movement patterns (Lédée et al. 2016) and the spatial structure of metapopulations (Dubois et al. 2016; Holstein et al. 2014); connecting biodiversity to ecosystem functioning (Creamer et al. 2016); identifying keystone species (Borrett 2013; Zhao et al. 2016); and using social network theory in studies of animal behavior (Croft et al. 2004; Fletcher et al. 2013; Krause et al. 2003; Sih et al. 2009). Further, ideas and concepts from network ecology are being applied to investigate the sustainability of urban and industrial systems (Fang et al. 2014; Layton et al. 2016; Xia et al. 2016) and elements of the food-energy-water nexus (Wang and Chen 2016; Yang and Chen 2016).

**Figure 1:**
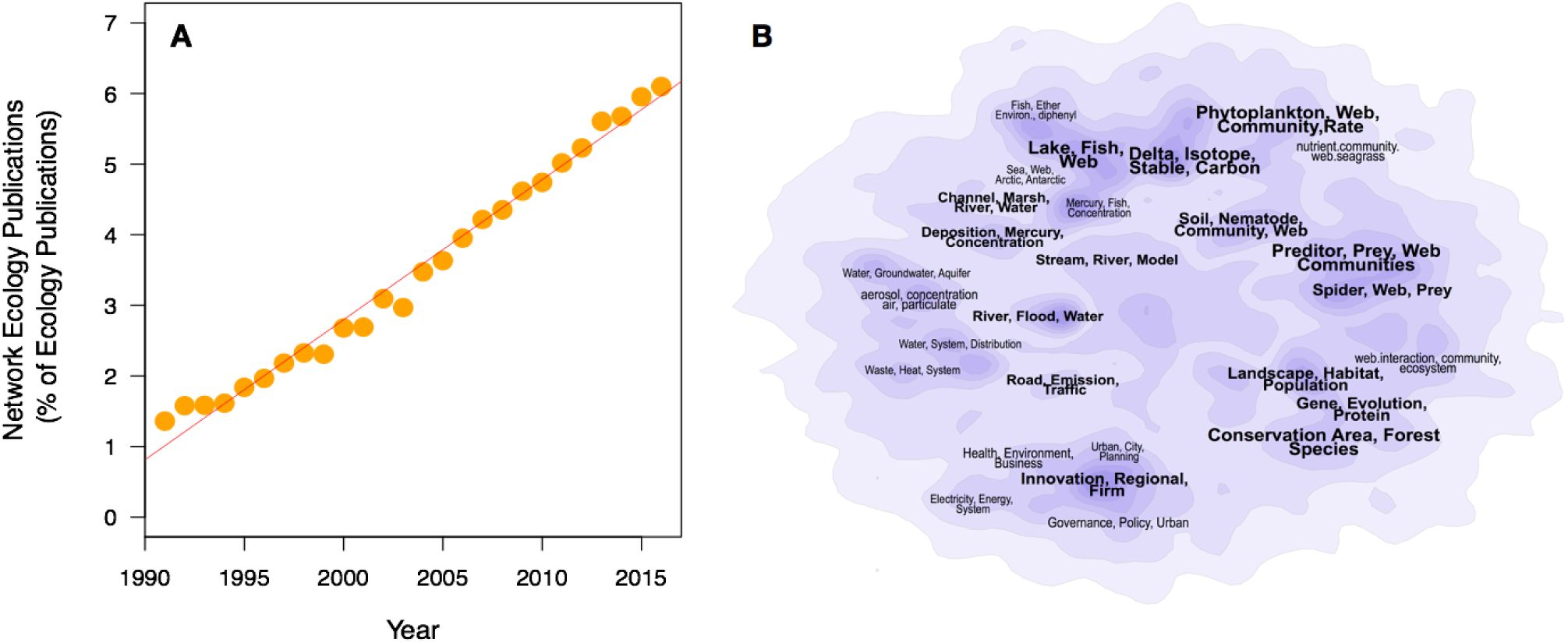
Although systems thinking has been a part of ecology since at least the work of Darwin, network ecology has grown rapidly since the turn of the last century but has been developing in isolated sub-fields. (A) Plot showing the increase in “network ecology” keywords in the literature from 1991 to current (updated using search developed by Borrett et al. (2014)). (B) Contour plot of common topics in network ecology with peaks indicating clusters of related topics. The regions are labeled with the most common terms found in the clusters. From Borrett et al. (2014), reproduced with permission.

Over the past 15 years, re-occurring themes for moving network ecology forward have emerged from reviews, perspectives, and syntheses (e.g., Bascompte 2010; Borrett et al. 2014; Poisot et al. 2015; Proulx et al. 2005). In this paper, we examine areas where the network approach is being applied to address important ecological questions and identify both challenges and opportunities for advancing the field. Among these are the need for shifting the focus toward mechanisms rather than observations, and increasing the resolution (e.g., individuals or traits as nodes and weighted edges of different interaction types) and replication of network models across different ecosystems and time (Ings et al. 2009; Poisot et al. 2016*b*; Woodward et al. 2010). After a brief primer of key concepts from network ecology, we discuss the following topics as they relate to these issues: the proliferation of terminology for ecological metrics with the increasing application of network methods; fully exploring the underlying assumptions of models of mechanistic processes for generating network structure; and the need for improved sharing and reproducibility of ecological network research and models. Although these topics are not new, the combination of the influx of metrics and theory and rapid increases in the computational intensity of ecology are creating novel challenges. With respect to these issues, we discuss recent advances that should be explored as tools to aid in a more effective integration of network methods for synthesis across ecological sub-disciplines.

## A primer of ecological networks: models and metrics

Prior to the introduction of network methods in ecology, the primary way of studying interactions was limited to detailed studies of behaviors and traits of individual species important to interactions, or of relationships between tightly interacting pairs of species (Carmel et al. 2013). Some ecologists were advancing whole-system methods (Lindeman 1942; Odum 1957); however, quantifying interactions is costly, as compared to surveys of species abundances. This has created a significant barrier to studying interactions at the scale of entire communities, either at the scale of individuals or species pairs, because the number of interactions becomes intractable. For instance, even if one assumes that only pairwise interactions occur among *S* species, the number of possible pairs is *S*(*S* − 1) /2. Local assemblages of macrobes often have 10^1^ – 10^2^ species, and microbial diversity can easily exceed 10^3^ OTUs (Operational Taxonomic Units).

This complexity of ecological systems is one reason there is a long tradition in community ecology of studying interactions within small subsets of closely-related species (e.g. trophic guilds) and using dimensionality reducing methods based on multivariate, correlative approaches (Legendre et al. 2012). While some approaches to studying subsets of species incorporate the underlying pattern of direct and indirect links (e. g., modules, (*sensu* Holt 1997; Holt and Hoopes 2005), the majority do not. Such limitations repeatedly have led to calls for the application of “network thinking” to ecological questions (e.g., Golubski et al. 2016; Ings et al. 2009; Jacoby and Freeman 2016; Patten and Witkamp 1967; Proulx et al. 2005; QUINTESSENCE Consortium et al. 2016; Urban and Keitt 2001). There are now many resources for learning about network ecology and network theory in general, and we point the reader in the direction of excellent reviews in this area (Bascompte and Jordano 2007; Borrett et al. 2012; Brandes et al. 2013; Ings et al. 2009; Proulx et al. 2005) and more comprehensive introductions (Brandes and Erlebach 2005; Estrada 2015; Newman 2010).

Network ecology employs network theory to quantify the structure of ecological interactions. All networks consist of sets of interacting nodes (e.g. species, non-living nutrient pools, habitat patches) whose relationships are represented by edges (e.g. nutrient or energy transfers, pollination, movement of individuals). Conceptually, a network is a set of things or objects with connections among them. Stated mathematically, a network is a generic relational-model comprised of a set of objects represented by nodes or vertices (*N*) and a set of edges (*E*) that map one or more relationships among the nodes, *G* = (*N, E*). A canonical ecological example of a network is a food-web diagram, in which the nodes represent species, groups of species, or non-living resources, and the *edges* map the relationship who-*eats*-whom.

The analysis of networks is inherently hierarchical, ranging from the entire network down to individual nodes and edges. Depending on the characteristics and level of detail of the information provided for a given model, there is a large number of network analyses and metrics that can be used to characterize the system at multiple levels (similar to Hines and Borrett, 2014; Wasserman and Faust, 1994), including: (1) the whole network level (i.e., the entire network), (2) the sub-network level (i.e., groups of two or more nodes and their edges), and (3) the individual node or edge level (Fig. 2).

**Figure 2:**
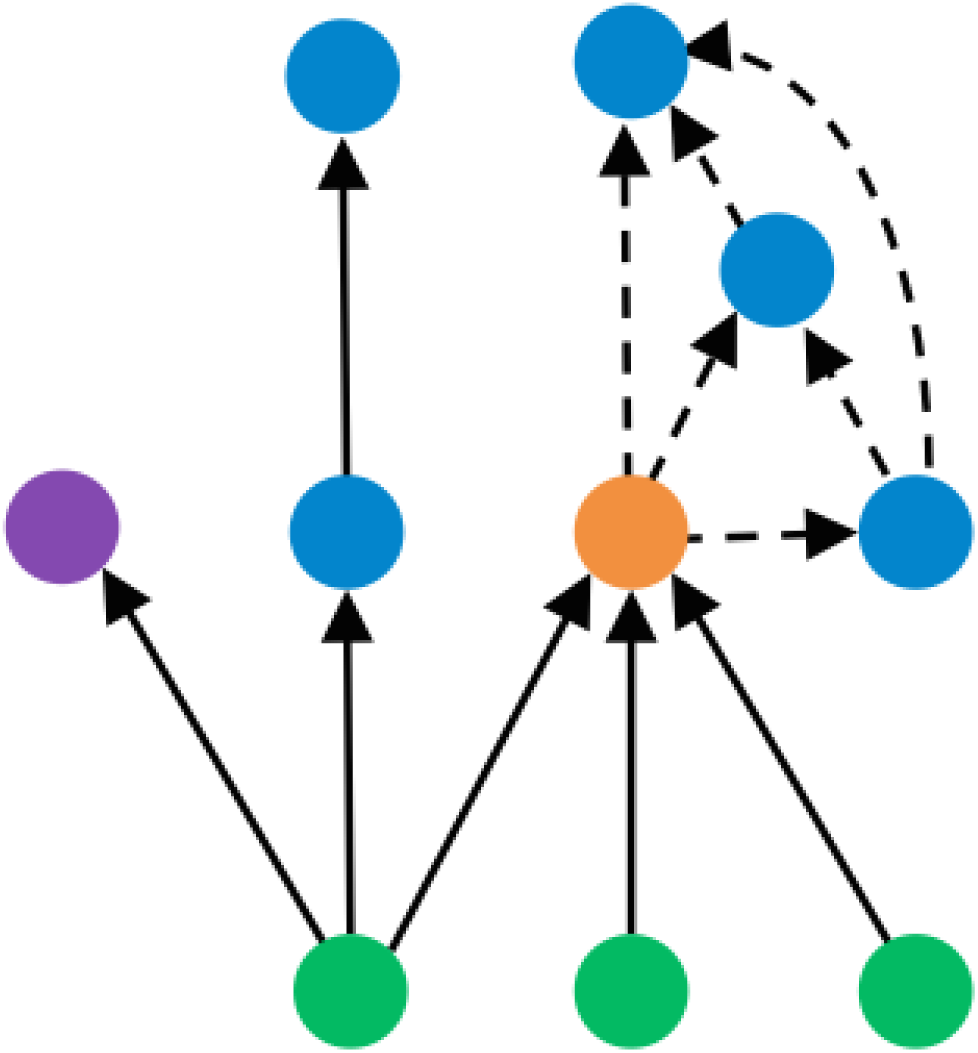
Hypothetical unweighted, directed network showing examples of the four classes of network metrics. *Node Level:* the purple node exhibits low centrality while the orange node exhibits high centrality. *Group or Sub-Network Level:* the blue nodes connected with dashed edges shows a module. *Global or Whole Network Level:* using the edges of all nodes we can measure the connectance of the entire network (*c* = *edges/nodes*^2^ = 0.12).

Network-level metrics integrate information over the entire set of nodes and edges. For example, the number of nodes (e.g., the species richness of a food web) and the density of connections or connectance are both network-level statistics used to describes the overall complexity of a network and have been investigated by ecologists for over 40 years (Allesina and Tang 2012; May 1972).

Sub-network level analyses focus on identifying specific subsets of nodes and edges. There are a variety of groups that have different names (e.g., module, motif, cluster, clique, environ) and different methods for measurement. Sub-networks often represent more tractable and meaningful units of study than individual nodes and edges on the one hand or entire networks on the other. For example, in landscape and population ecology, the preferential movement of individuals and genes (edges) between habitat patches (nodes) has implications for conservation of populations and the design of preserves (Calabrese and Fagan 2004; Fletcher et al. 2013; Holt and Hoopes 2005). Also, both nodes and edges can be divided into classes. An example of this is the bipartite graph, in which interactions occur primarily between, rather than within, each class or “part” of the community. A bipartite network has only two classes of nodes, such as in a pollination network in which the community is divided into plants being pollinated and insects that do the pollination (Petanidou et al. 2008). In this network, edges representing pollination visits can only map between two nodes in the different classes.

Metrics at the individual node or edge level quantify differences in relative importance. Whether we are interested in an individual or species that transmits disease, species whose removal will result in secondary extinctions, or key habitat patches that connect fragmented landscapes, identifying important nodes is a critical component of network analysis. Another type of node or edge-level metric classifies nodes or edges according to their roles within a network. This classification can use information from differing levels. Additionally, nodes and edges can have variable characteristics. Edges can be weighted and they can map a directed relationship (as opposed to a symmetric or undirected relationship). For example, in ecosystem networks, the edges show the directed movement of energy or nutrients from one node to another by some process like feeding, and the edge weight can indicate the amount of energy or mass in the transaction (Baird and Ulanowicz 1989; Dame and Patten 1981). Nodes also can be weighted (e.g., size of individual, population size, biomass of a given species). Lastly, network models are flexible enough to accommodate variation in edge types and relationships among edges (e.g., hypergraphs), but analysis of these more complicated models is challenging and has only begun to be applied in ecology (e.g., Golubski et al. 2016).

## Resolving network metrics

The application of network theory defines an explicit mathematical formalism that provides a potentially unifying set of terms for ecology and its inter-disciplinary applications (QUINTESSENCE Consortium et al. 2016). Ironically, the development of ecological network metrics has had an opposing affect. One reason for this is that introductions have occurred in multiple sub-disciplinary branches (Fig. 1B) (Blüthgen 2010; Borrett et al. 2014; Carmel et al. 2013). Having separate research trajectories can facilitate rapid development of ideas and the process of integration can lead to novel insights (Hodges 2008). At the same time, these innovations in network ecology have come at the cost of the “rediscovery” of the same network metrics and subsequent description of them with new terms. This has led to different metrics with similar purposes existing in separate areas of ecology (Table 1).

**Table 1:** Ecological network metric summary and classification. Level indicates the hierarchy of the metric (W = Whole network, G = Group or sub-network, N = Node). The Sub-disciplines include ’General’ network theory, ’Community’ ecology, ’Food-web’ and ’Ecosystem’ ecology. Also available at https://figshare.com/s/1bf1a7e0a6ee3ac97a4b.

| Sub.discipline | Level | Metric | Concept | Reference |
| --- | --- | --- | --- | --- |
| General | W | Density | The proportion of possible edges that are actually associated with nodes; called Connectance in Food Web ecology. |  |
| General | N | Centrality | Multiple ways to characterize the relative importance of nodes. | Wasserman and Faust (1994) |
| General | N | Degree | Number of edges connected to a given node, which is a type of local centrality. |  |
| General | N | Eigenvector Centrality | Global centrality metric based on number of walks that travel through a node | Bonacich (1987) |
| General | W | Centrality Distribution | Shape of the frequency distribution of edges among nodes. | Barabási and Albert (1999); Dunne et al. (2002) |
| General | W | Centralization | The concentration (versus evenness) of centrality among the nodes. | Freeman (1979) |
| General | W | Graph diameter | The longest path between any two nodes in a graph. | Barabási et al. (2000); Urban and Keitt (2001) |
| General | W | Modularity | Degree to which edges are distributed within rather than between distinct sets of nodes. | Newman (2010) |
| General | G | Motifs | Small sets of nodes with similar distributions of edges. | Milo et al. (2002) |
| General | W | Link density | Average number of edges per node. | Martinez (1992) |
| Community | N | Temperature | Measures the nestedness of a bipartite network. | Ulrich and Gotelli (2007) |
| Community | W | Co-occurrence | Degree of overlapping spatial or temporal distributions of species relative to a null model. | Gotelli (2000) |
| Community | N | Indicator Species | The degree to which the abundance of a taxonomic group responds to an environmental gradient. |  |
| Community | W | Nestedness | The degree to which interactions can be arranged into subsets of the larger community |  |
| Community | W | Evenness | Deviation of the distribution of observed abundances relative to an even distribution among taxonomic groups in a community |  |
| Community | W | Diversity | Distribution of abundances among taxonomic groups in an observed community |  |
| Community | W | Richness | The number of taxonomic groups in a community |  |
| Community | W | Stability | The change in the abundances of taxonomic groups across a set of observations |  |
| Food-Web | N | Removal Importance | The degree to which removal of a compartment or species produces subsequent removals in the ecosystem. | Borrvall et al. (2000); Dunne et al. (2002); Eklöf and Ebenman (2006); Solé and Montoya (2001) |
| General | N | Connectance | Proportion of realized out of possible edges | Pimm (1982); Vermaat et al. (2009) |
| Food-Web | G | Food-chain length | The number of feeding relationships among a set of compartments in a food-web. | Post et al. (2000); Ulanowicz et al. (2014) |
| Ecosystem | W | Finn cycling index | Degree to which matter or energy passes through the same set of compartments. | Finn (1980) |
| Ecosystem | G | Environ | The sub-network of the probability of movement of energy or matter among compartments generated by a single unit of input (output) into a selected node. | Patten (1978); Patten and Auble (1981) |
| Ecosystem | N | Throughflow | Amount of energy or matter passing into or out of a node | Finn (1976) |
| Ecosystem | N | Throughflow Centrality | The proportion of energy or matter that passes through a given compartment in an ecosystem. | Borrett (2013) |
| General | G | Chain Length | Number of edges between two nodes in a group |  |
| Food-Web | G | Average Path Length | The average number of times a unit of matter or energy travels from one compartment to another before exiting the ecosystem | Finn (1976) |
| Ecosystem | W | Pathway Proliferation | Rate of increase in the number of edges between nodes with increasing path length | Borrett et al. (2007) |
| Ecosystem | W | Ascendency | Measures the average similarity in matter or energy flows among compartments in an ecosystem. | Ulanowicz (1986) |
| Food-Web | N | Trophic Level | Ordinal classification of a compartment or taxonomic group based on the relative position in the ecosystem. | Allesina and Pascual (2009); Fath (2004); Williams et al. (2002) |

Ecological studies using network approaches draw from a deep well of general network theory (Newman 2003, 2006; Strogatz 2001). Ecologists broadly use network concepts, techniques, and tools to: (1) characterize the system organization (Borrett 2013; Croft et al. 2004; Ulanowicz 1986); (2) investigate the consequences of the network organization (Borrett et al. 2006; Dunne et al. 2002; Grilli et al. 2016); and (3) identify the processes or mechanisms that might generate the observed patterns (Allesina and Pascual 2008; Fath et al. 2007; Guimarães et al. 2007; Poisot et al. 2016*b*; Ulanowicz et al. 2014; Williams and Martinez 2000). The unnecessary proliferation of network metrics is exemplified by “connectance” (*C*), which is used by food-web ecologists to mean the ratio of the number of edges in the network divided by the total number of possible edges. Elsewhere in the network science literature, this measurement is referred to as network density (Newman et al. 2001). As an-other example, what ecosystem ecologists have described as “average path length” (total system through-flow divided by the total system input) (Finn 1976) also has been called network aggradation (Jørgensen et al. 2000). In economics, average path length is known as the multiplier effect (Samuelson 1948).

Another kind of redundancy is the creation and use of multiple statistics that measure the same or very similar network aspects. A clear example of this is inherent in the proliferation of centrality measures to indicate node or edge importance. Network scientists have shown that many centrality metrics are correlated (Jordán et al. 2007; Newman 2006; Valente et al. 2008). Likewise, Borrett and Osidele (2007) found that nine commonly reported ecosystem network analysis metrics covaried in 90 plausible parameterizations of a model of phosphorus biogeochemical cycling for Lake Lanier, GA, but that all these metrics were associated strongly with only two underlying factors. However, even a perfect correlation does not mean that two metrics have identical properties, and they still may diverge in different models. Therefore, it is important to have mathematically based comparisons of metrics (Borgatti and Everett 2006; Borrett 2013; Kazanci and Ma 2015; Ludovisi and Scharler 2017). It is incumbent on network ecologists to establish clearly the independence and uniqueness of the descriptive metrics used.

From the perspective of the broader field of ecology, the proliferation of concepts, terms, and metrics is not a new issue (e.g., Ellison et al. 2005; Tansley 1935). Ecologists have a long history of using network concepts and related models in multiple subdomains (e.g., metapopulations, matrix population models, community co-occurrence models, ecosystems) without fully recognizing or capitalizing on the similarities of the underlying models. Each subdomain has constructed its own concepts and methods (occasionally borrowing from other areas), and established its own jargon that impedes scientific development. Previous suggestions for solving this issue have focused on maintaining an historical perspective of ecology (Graham and Dayton 2002); Blüthgen et al. (2008) is an excellent example of how this can be done through peer-reviewed literature.

One possible approach that would go beyond such a diffuse, literature-centered approach would be to develop a formal ontology of concepts and metrics. An on-tology is a a set of related terms that are formally defined and supported by assertions (Bard and Rhee 2004). An ontology therefore provides a framework for developing concepts within a discipline and presents the opportunity for more efficient synthesis across disciplinary boundaries. The concept of an ontology is not new, but more rapid sharing of ontologies and their collaborative development have been enabled by the Internet. For example, the Open Biological Ontologies (OBO, http://www.obofoundry.org) supports the creation and sharing of ontologies over the web. Currently, there is no OBO for a “network ecology metric” ontology, and as far as we are aware, ontologies have yet to be explored or developed for network metrics.

The OBO could provide a platform for harmonizing ecological network metrics, terms, and concepts. Key obstacles to such harmonization include a requirement that network ecologists work within a common framework, and the need for an individual or leadership team to periodically curate the ontology based on new developments in the field. In determining the best course of action, network ecologists could follow the example of how similar OBO projects have been managed in the past. The *FOODON* food role ontology project (http://www.obofoundry.org/ontology/foodon.html) contains information about “materials in natural ecosystems and food webs as well as human-centric categorization and handling of food.” It could serve as an example or even the basis of a ecological network metric ontology.

## Benchmarking: Trusting our models of mechanisms

Inferences about processes in ecological systems have relied in part on the application of simulation models that generate matrices with predictable properties. As discussed in the previous section, the proliferation of network metrics points to the need for the investigation and comparison of how these metrics will behave in the context of different modeling algorithms. Once a metric or algorithm has been chosen, it is tempting apply them widely to empirical systems to detect patterns, but before research proceeds, a process of “benchmarking” with artificial matrices that have predefined amounts of structure and randomness should be used to examine the behavior of the algorithms and the metrics that are applied to them.

Benchmarking of ecological models developed from null model analysis in community ecology (Atmar and Patterson 1993; Connor and Simberloff 1979; Gotelli and Ulrich 2012). Null models are specific examples of randomization or Monte Carlo tests (Manly 2007) that estimate a frequentist *P* value, the tail probability of obtaining the value of some metric if the null hypothesis were true (Gotelli and Graves 1996). The aim of a null model is to determine if the structure of an observed ecological pattern in space or time is incongruous with what would be expected given the absence of a causal mechanism. A metric of structure calculated for a single empirical data set is compared to the distribution of the same metric calculated for a collection of a large number of randomizations of the empirical data set. The data are typically randomized by reshuffling some elements while holding other elements constant to incorporate realistic constraints. Comparison with a suite of null models in which different constraints are systematically imposed or relaxed may provide a better understanding of the factors that contribute most to patterns in the network (see Box 1). However, the devil remains in the details and there are also a variety of ways to randomize data and impose constraints to construct a useful null model. If the null model is too simplistic, such as a model in which edges and nodes are sampled with uniform probability, it will always be rejected and provide little insight into important ecological patterns, regardless of what metric is used. At the other extreme, if the null model incorporates too many constraints from the data, it will be difficult or impossible to reject the null hypothesis, even though the network may contain interesting structure.

#### Boxes

*Box 1. Benchmarking Ecological Models* The most basic test is to feed the algorithm a set of “random” matrices to make sure that the frequency of statistically significant results is no greater than 5%. Otherwise, the algorithm is vulnerable to a Type I statistical error (incorrectly rejecting a true null hypothesis). However, specifying a matrix produced by random sampling errors is not so easy. By definition, if a null model algorithm is used to generate the random matrices, then no more than 5% of them should be statistically significant (unless there were programming errors). For binary matrices, two log-normal distributions can be used to generate realistic heterogeneity in row and column totals, while still maintaining additive effects for cell occurrence probabilities (Ulrich and Gotelli 2010). “Structured” matrices are needed to test for Type II errors (incorrectly accepting a false null hypothesis), and these require a careful consideration of exactly what sort of pattern or mechanism the test is designed to reveal. One approach is to begin with a perfectly structured matrix, such as one derived from a mechanistic model for generating network structure, contaminate it with increasing amounts of stochastic noise, and test for the statistical pattern at each step (Gotelli 2000). A plot of the *P* value versus the added noise should reveal an increasing curve, and will indicate the signal-to-noise ratio below which the test cannot distinguish the pattern from randomness. Alternatively, one can begin with a purely random matrix but embed in it a non-random substructure, such as a matrix clique or a node with extreme centrality. The size, density, and other attributes of this matrix can be manipulated to see whether the test can still detect the presence of the embedded structure (Gotelli et al. 2010). Because all null model tests (and all frequentist statistics) are affected by sample size and data structure, these benchmark tests can be tailored to the attributes of the empirical data structures for better focus and improved inference.

Even simple randomization algorithms may require further filters to ensure that random matrices retain a number of desirable network properties. For example, Dunne et al. (2002) created random food-web matrices with constant species richness and connectance, but they discarded webs with unconnected nodes and subwebs because these topologies were not observed in the empirical webs. A “stub reconstruction” algorithm builds a topology that is constrained to the observed number of edges per node (Newman et al. 2001). Each node is assigned the correct number of edges, and then nodes are successively and randomly paired to create a growing network. However, this algorithm also generates multiple edges between the same two nodes, which must be discarded or otherwise accounted for. Maslov and Sneppen (2002) use a “local re-wiring algorithm” that preserves the number of connections for every node by swapping edges randomly between different pairs of nodes. This algorithm is closely analogous to the swap algorithm used in species co-occurrence analyses that preserves the row and column totals of the original matrix (Connor and Simberloff 1979). The more constraints that are added to the algorithm, the less likely it is that simple sampling processes can account for patterns in the data. However, some constraints, such as connectivity or matrix density, may inadvertently “smuggle in” the very processes they are designed to detect. This can lead to the so-called “Narcissus” effect (Colwell and Winkler 1984). Finding the correct balance between realistic constraints and statistical power is not easy (Gotelli et al. 2012), and there are many potential algorithms that reasonably could be used, even for simple binary matrices (Gotelli 2000).

In network theory, the Erdos-Renyi (ER, (Erdös and Rényi 1959)) model is a now-classic example of a model used to generate networks via a random process for creating matrix structure. The ER model is a random graph that starts with an *N* × *N* adjacency matrix of nodes and assigns to it *K* edges between randomly chosen pairs of nodes. The ER model has been applied in ecology to address questions about the relationship between stability and complexity (May 1972) and the structure of genetic networks (Kauffman et al. 2003). For example, randomized networks have been used to link motifs (Milo et al. 2002) to network assembly (Baiser et al. 2016), stability (Allesina and Pascual 2008; Borrelli et al. 2015), and persistence in food webs (Stouffer and Bascompte 2010).

In addition to the random matrix approaches of null and ER models, there are other, more complex algorithms that are used to generate structured matrices. Perhaps one of the best known in network theory is the Barabasi-Albert (BA, Barabási and Albert 1999) model, which adds nodes and edges to a growing network with a greater probability of adding edges to nodes with a higher degree. The BA algorithm is similar to ecological network algorithms that generate non-random structure, because of either direct influence or similar processes operating in systems of interest. Some of these models include processes of “preferential attachment” in which organisms tend to interact with the same, common species. Food-web modeling algorithms also have been developed that use a trait-based approach (e.g., Allesina and Pascual 2009), consumer-resource models (Yodzis and Innes 1992), niches (Williams and Martinez 2000), cyber-ecosystem algorithms (Fath 2004), and cascade models (Allesina and Pascual 2009; Allesina and Tang 2012; Cohen and Łuczak 1992).

The statistical behavior of some models and metrics can be understood analytically. For example, the networks generated by the BA algorithm display degree distributions that approximate a power-law distribution, like many real-world “scale-free” networks (Albert et al. 2002). If the network is sparse (i.e., (*K* ≪ *N*^2^)), the degree distribution of the network should follow a Poisson distribution. However, as new models and metrics are introduced, new benchmarking should be done and compared to previous results. Newman et al. (2016) is one example of how benchmarking can be used for investigating processes operating on ecological networks. Ludovisi and Scharler (2017) advocate the same approach for the analysis of network models in general. The `benchmark` (Eugster and Leisch 2008) package in R (R Core Team 2017) is a general algorithm-testing software package that provides a useful starting point.

## Reproducibility: Open-data, Open-source and Provenance

As analyses of network models increase in computational intensity, there is a concomitant increase in the need for new tools to track and share key computational details. This need is compounded when models incorporate data from multiple sources or analyses involve random processes. The combination of the volume of data and computational intensity of studies of ecological networks further increases the burden on ecologists to provide information needed to adequately reproduce datasets, analyses, and results. As the sharing and reproducibility of scientific studies are both essential for advances to have lasting impact, finding easier, faster, and generally more convenient ways to record and report relevant information for ecological network studies is imperative for advancing the field.

Sharing data and open-source code have become established in ecology, and network ecologists are now producing more network models and data (e.g., Fig. 1A). These include not only ecological interaction networks, but also an influx of other relevant networks, including ecological genomic networks generated by next-generation, high-throughput sequencing technologies (Langfelder and Horvath 2008; Zinkgraf et al. 2017). There are now multiple web-accessible scientific databases (e.g., NCBI, Data Dryad, Dataverse) and at least four databases have been constructed specifically to curate ecological network data: including “Kelpforest” (Beas-Luna et al. 2014), “The Web of Life” (Fortuna et al. 2014), “Mangal” ecological network database (Poisot et al. 2015) and the “Interaction Web Database” (https://www.nceas.ucsb.edu/interactionweb/resources.html).

The increase in ecological network data is linked to an increasing rate of shared analytical code and other open-source software. It is now commonplace for ecologists to have a working knowledge of one or more programming languages, such as R, Python, SAS, MatLab, Mathematica, or SPSS. Multiple software packages exist for doing ecological analyses, including ecological network analyses. In addition to the general network analysis packages available in R, there are at least two packages aimed specifically at ecological network analysis: `bipartite` and `enaR`. The former provides functions drawn largely from community ecology (Dormann et al. 2009), whereas the latter provides a suite of algorithms developed in the ecosystem network analysis literature (Borrett and Lau 2014; Lau et al. 2015).

Although, ecology has long had a culture of keeping records of important research details, such as field and lab notebooks, these practices put all of the burden of recording “metadata” on the researcher. Manual record-keeping methods, even when conforming to metadata standards (e.g., EML, see Boose et al. 2007), do not take advantage of the power of the computational environment. Data-provenance methods aim to provide a means to collect formalized information about computational processes, ideally in a way that aids the reproducibility of studies with minimal impact on the day-to-day activities of researchers (Boose et al. 2007). These techniques have been applied in other areas of research and could provide an effective means for documenting the source and processing of data from the raw state into a model (Boose and Lerner 2017).

The reproducibility of scientific studies is imperative for advances to have lasting impact through the independent verification of results. Although this has been an ongoing topic of discussion in ecology (Ellison 2010; Parker et al. 2016), the need was highlighted by a recent survey finding issues with reproduction of studies across many scientific disciplines (Baker 2016). There is significant motivation from within the ecological community to move toward providing detailed information about computational workflows for both repeatability and reproducibility, which includes repetition by the original investigator (Lowndes et al. 2017). It is also important in network ecology for data sources and methods for model construction be standardized and transparent, and that models be curated and shared (McNutt et al. 2016).

Collecting details, such as those enabled by data-provenance capture software, is one innovative way forward. These tools have been developing in the computer-science domain for decades; however, only recently have they gained a foothold in ecology (Boose et al. 2007; Ellison 2010) or the broader scientific community. Although there are many challenges in the development and application of data-provenance principles, multiple software packages do exist for collecting data provenance in the context of scientific investigations. Two provenance capture packages exist in R, the `recordr` package associated with the DataOne repository (Cao et al. 2016) and `RDataTracker` (Lerner and Boose 2014). In addition, although they do not collect formal data provenance, there are methods developed for “literate computing” that help to collect code along with details about the code and the intention of the analyses (e.g., the Jupyter notebook project: (Shen and Barabasi 2014)).

For ecological networks, there is software that captures the “data pedigree” of food-web models, but it does not capture data provenance. Data pedigree was initially implemented in the EcoPath food-web modeling package (Guesnet et al. 2015; Heymans et al. 2016) to define confidence intervals and precision estimates for network edges. It has been developed further to allow for the use of informative priors in Bayesian modeling of ecological networks. This is done by linking models to the literature sources from which estimates were derived, an approach that is similar to incorporating metadata information within databases of ecological networks. Although this approach focuses only on a subcomponent of provenance, this still is a promising way to address the issue that networks, network metrics, and simulation models used to analyze them commonly assume a lack of uncertainty (e.g., Borrett and Osidele 2007; Kauffman et al. 2003; Kones et al. 2009), and typically ignore inaccuracy in the empirical data (Ascough et al. 2008; Gregr and Chan 2014).

## Moving Forward

Development and application of new technologies (e.g., sequencing methods and computational, data-driven approaches) have the potential to increase both the abundance and quality of ecological networks. For the future development of network ecology, there is a pressing need not only to share data and code, but also to integrate and use the large amounts of information enabled by technological advances. For example, synthetic networks (i.e., merging network models from different studies Poisot et al. 2016*a*) are a promising new direction; however, the structural properties of synthetic networks and the behavior of network metrics applied to them will require careful investigation, including the application of systematic benchmarking. Multi-trophic networks provide a precedence for these studies to move forward, but synthesizing models from across many different sources produces new challenges for developing and benchmarking metrics, as well as an opportunity for new technologies, like data provenance, to help establish better connections among studies and researchers.

The burgeoning of “open” culture in the sciences (Hampton et al. 2014) also has the potential to serve as a resource for models and a clearinghouse for resolving the validity of metrics, models, and algorithms. First, because code is openly shared, functions used to calculate metrics are open for inspection and, if coded and documented clearly using software “best-practices” (e.g., Noble 2009; Visser et al. 2015), the code provides a transparent documentation of how a metric is implemented and its computational similarity to other metrics. Second, enabled by the ability to write their own functions and code, researchers can do numerical investigations of the similarities among metrics. Through comparison of metrics calculated on the same or similar network models, a researcher could at least argue, for a given set of models, that two or more metrics produce similar results. Third, data provenance provides a useful tool to aide in the dissemination and synthesis of network models and increases the reproducibility of ecological network studies, including those documenting new metrics and benchmarking those metrics and associated algorithms for generating or analyzing empirical models. Last, as with data provenance, formalizing ecological network metrics and concepts requires a mathematically rigorous foundation that is developed by the community of researchers working along parallel lines of inquiry. Whether this is done through an ontological approach or some other formalized “clearing-house,” an open process of exchange that integrates multiple perspectives is essential to prevent the rapid dilution of concepts in ecological network research as these concepts continue to proliferate, develop and evolve.

Over half a century ago, Robert MacArthur published his first paper on the relationship between diversity and stability, initiating multiple research trajectories that have now become the mainstay of many ecological research programs (MacArthur 1955). The theory that MacArthur applied was based on flows of energy through networks of interacting species. Thus, network theory is at the roots of one of the most widely studied topics in ecology and is now a part of the broader context of integration across many scientific disciplines that is aimed at consilience of theory (Wilson 1999). The synthesis of ecological concepts through the mathematically rigorous “lingua franca” of network terminology has the potential to unify theories across disciplines. As with previous concepts (e.g., keystone species, foundation species, ecosystem engineer), greater clarity and less redundancy will come about as network methods are used more commonly and researchers compare the mathematical and computational underpinnings of the metrics that they are using. With the increased use of these approaches, the network concept has and will continue to serve as a common model that transcends disciplines and has the potential to serve as an inroad for new approaches. With thoughtful dialogue across sub-disciplines and among research groups, further infusion of network theory and methods will continue to advance ecology.

## Acknowledgments

This work was supported by the US National Science Foundation under grant SSI-1450277 End-to-End Provenance.

## Author contributions statement

All authors contributed to the conception, writing and review of the manuscript.

